# LY6E Restricts the Entry of Human Coronaviruses, including the currently pandemic SARS-CoV-2

**DOI:** 10.1101/2020.04.02.021469

**Authors:** Xuesen Zhao, Shuangli Zheng, Danying Chen, Mei Zheng, Xinglin Li, Guoli Li, Hanxin Lin, Jinhong Chang, Hui Zeng, Ju-Tao Guo

## Abstract

C3A is a sub-clone of human hepatoblastoma HepG2 cell line with the strong contact inhibition of growth. We fortuitously found that C3A was more susceptible to human coronavirus HCoV-OC43 infection than HepG2, which was attributed to the increased efficiency of virus entry into C3A cells. In an effort to search for the host cellular protein(s) mediating the differential susceptibility of the two cell lines to HCoV-OC43 infection, we found that ADAP2, GILT and LY6E, three cellular proteins with known activity of interfering virus entry, expressed at significantly higher levels in HepG2 cells. Functional analyses revealed that ectopic expression of LY6E, but not GILT or ADAP2, in HEK 293 cells inhibited the entry of HCoV-OC43. While overexpression of LY6E in C3A and A549 cells efficiently inhibited the infection of HCoV-OC43, knockdown of LY6E expression in HepG2 significantly increased its susceptibility to HCoV-OC43 infection. Moreover, we found that LY6E also efficiently restricted the entry mediated by the envelope spike proteins of other human coronaviruses, including the currently pandemic SARS-CoV-2. Interestingly, overexpression of serine protease TMPRSS2 or amphotericin treatment significantly neutralized the IFITM3 restriction of human coronavirus entry, but did not compromise the effect of LY6E on the entry of human coronaviruses. The work reported herein thus demonstrates that LY6E is a critical antiviral immune effector that controls CoV infection and pathogenesis *via* a distinct mechanism.

**Importance:** Virus entry into host cells is one of the key determinants of host range and cell tropism and is subjected to the control by host innate and adaptive immune responses. In the last decade, several interferon inducible cellular proteins, including IFITMs, GILT, ADAP2, 25CH and LY6E, had been identified to modulate the infectious entry of a variety of viruses. Particularly, LY6E was recently identified as host factors to facilitate the entry of several human pathogenic viruses, including human immunodeficiency virus, influenza A virus and yellow fever virus. Identification of LY6E as a potent restriction factor of coronaviruses expands the biological function of LY6E and sheds new light on the immunopathogenesis of human coronavirus infection.

## INTRODUCTION

Coronaviruses (CoV) are a large family of enveloped positive-strand RNA viruses with broad host ranges and tissue tropism (1, 2). While four human CoVs, including HCoV-229E, HCoV-OC43, HCoV-NL63 and HCoV-HKU1, cause mild upper respiratory tract infections, three zoonotic CoVs have crossed species barriers to infect humans since 2002 and cause severe acute respiratory syndrome (SARS) (3, 4), Middle East respiratory syndrome (MERS) (5, 6) and coronaviral disease-19 (COVID-19) (7, 8), with the mortality rate of 10%, 30% and 1 to 2%, respectively (9, 10). No vaccine or antiviral drug is currently available to prevent CoV infection or treat the infected individuals. The cross-species transmission of zoonotic CoVs presents a continuous threat to global human health (11, 12). Therefore, understanding the mechanism of CoV infection and pathogenesis is important for the development of vaccines and antiviral agents to control the current COVID-19 pandemics and prevent future zoonotic CoV threats.

CoV entry into host cells, a process to deliver viral nucleocapsids cross the plasma membrane barrier into the cytoplasm, is the key determinant of virus host range and plays a critical role in zoonotic CoV cross-species transmission (2, 13). The entry process begins by the binding of viruses to their specific receptor on the plasma membrane, which triggers endocytosis to internalize the viruses into the endocytic vesicles. The cleavage of viral envelope spike proteins by endocytic proteases and/or endosomal acidification triggers the conformation change of spike protein to induce the fusion of viral envelope with endocytic membrane and release nucleocapsids into the cytoplasm to initiate viral protein synthesis and RNA replication. While angiotensin-converting enzyme 2 (ACE2) is the *bona fide* receptor for SARS-CoV, SARS-CoV-2 and HCoV-NL63 (14-16), MERS-CoV and HCoV-229E use dipeptidyl peptidase-4 (DPP4) and CD13 (also known as aminopeptidase N) as their receptor, respectively (17, 18). However, HCoV-OC43 and HCoV-HKU1 bind to 9-Oacetylated sialic acids *via* a conserved receptor-binding site in spike protein domain A to initiate the infection of target cells (19). As the key determinant of cell tropism, host range, and pathogenesis, CoV entry is primarily controlled by interactions between the spike envelope glycoprotein and host cell receptor as well as the susceptibility of spike glycoprotein to protease cleavage and/or acid-induced activation of membrane fusion (20, 21). For instance, SARS-CoV can use ACE2 orthologs of different animal species as receptors (22-26) and the efficiency of these ACE2 orthologs to mediate SARS-CoV cell entry is consistent with the susceptibility of these animals to SARS-CoV infection (27-30). In addition, expression of endosomal cathepsins, cell surface transmembrane proteases (TMPRSS), furin, and trypsin differentially modulates the entry of different human CoVs (31-35).

Interferons (IFNs) are the primary antiviral cytokines that mediate innate and adaptive immune control of virus infection by inducing hundreds of genes, many of which encode antiviral effectors (36). In the last decades, several IFN-inducible proteins, including three IFN-induced transmembrane (IFITM) proteins (37), gamma-interferon-inducible lysosome/endosome-localized thiolreductase (GILT) (38), 25-Hydroxycholesterol hydrolase (25HC) (39), ArfGAP with dual pleckstrin homology (PH) domains 2 (ADAP2) (40) and lymphocyte antigen 6 family member E (LY6E) (41) had been identified to restrict or enhance the entry of a variety of viruses. Interestingly, while IFITM proteins inhibit the entry of all the other human CoVs, HCoV-OC43 hijacks human IFITM2 or IFITM3 as entry factors to facilitate its infection of host cells (42, 43). We also demonstrated recently that GILT suppresses the entry of SARS-CoV, but not other human CoVs (38). As reported herein, in our efforts to identify host factor(s) determining the differential susceptibility of two closed related human hepatoma cell lines to HCoV-OC43 infection, we found that LY6E potently suppresses the infectious entry of all the human CoVs, including the currently pandemic SARS-COV-2. Our study also revealed that unlikely IFITMs, LY6E inhibits CoV entry *via* a distinct mechanism.

## RESULTS

### C3A is more susceptible to HCoV-OC43 infection than its parental cell line HepG2

C3A is a sub-clone of HepG2 that was selected for strong contact inhibition of growth and high albumin production (44). Metabolically, C3A is more relevant to normal hepatocytes and has been used for development of bioartificial liver devices (45). Interestingly, we found that these two closely related cell lines drastically differ in their susceptibility to HCoV-OC43 infection (Fig. 1). Specifically, infection of the two cell lines with the virus at a MOI of 0.02, 0.2 and 1 resulted in approximately 75, 25 and 10 folds more infected cells in C3A cultures than in HepG2 cultures at 24 h post infection, respectively (Fig. 1A). Consistent with this finding, much higher levels of viral nucleocapsid protein (N) and RNA were detected in infected C3A cultures (Figs. 1B and C). Infected C3A cultures also produced approximately 20-fold more progeny viruses than HepG2 cultures did (Fig. 1D). To determine whether the differential susceptibility of the two hepatoma cell lines to HCoV-OC43 infection is due to a difference in virus entry or post-entry replication event, we compared the susceptibility of the two cell lines to lentiviral particles pseudotyped with envelope proteins of HCoV-OC43, influenza A virus (IAV), vesicular stomatitis virus (VSV) or Lassa fever virus (LASV). As shown in Fig 2, while pseudoviral particles of IAV (IAVpp), VSV (VSVpp) and LASV (LASVpp) infected the two cell lines with similar efficiency, the efficiency of HCoV-OC43pp infection in C3A cultures is approximately 50 folds higher than that in HepG2 cultures. These results clearly indicate that the differential susceptibility is attributed to the distinct ability of the two cell lines to support the infectious entry of HCoV-OC43.

**Fig. 1.**
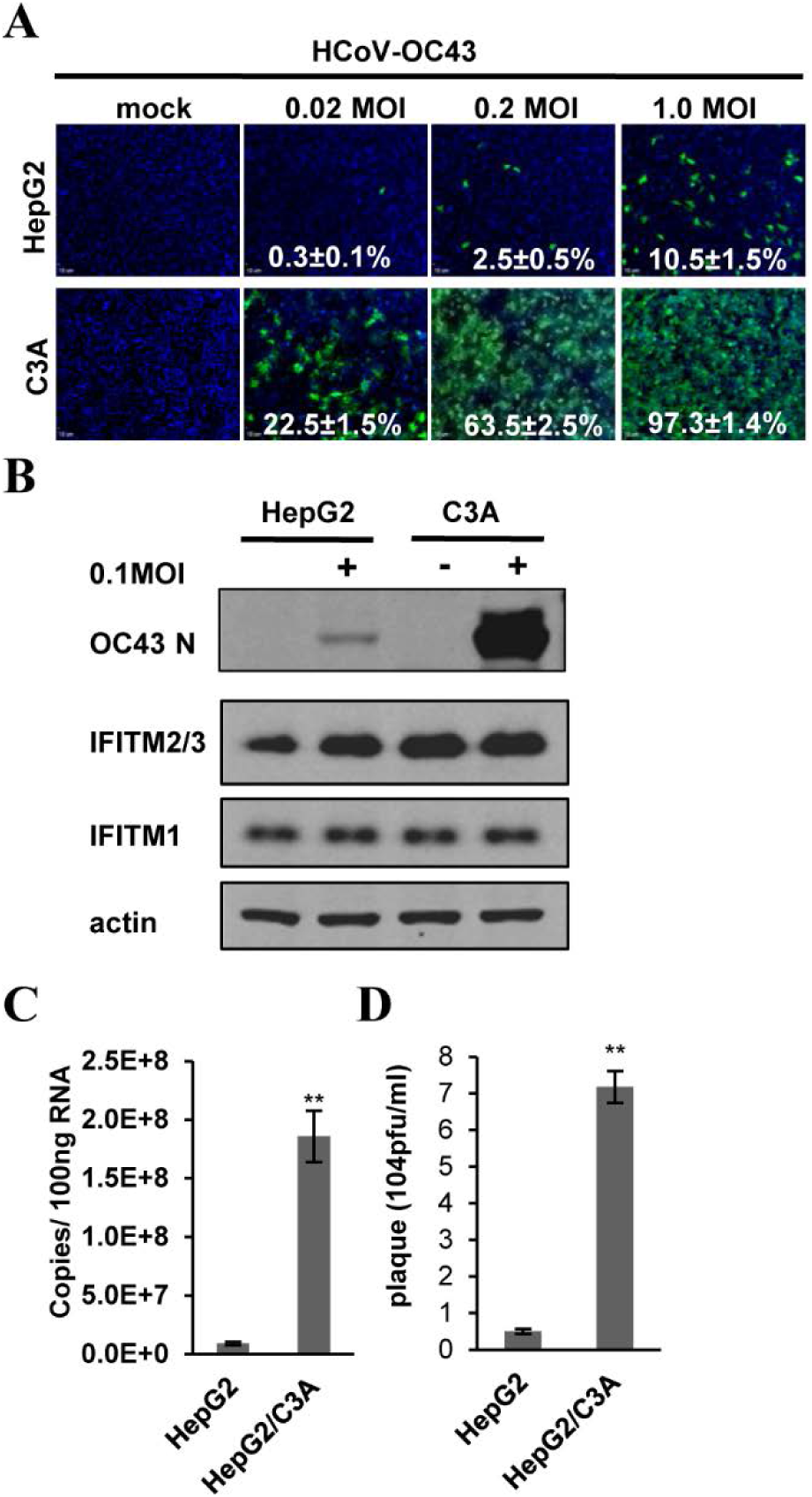
C3A cells are more susceptible to HCoV-OC43 infection than HepG2 cells. HepG2 and C3A cells were mock-infected or infected with HCoV-OC43 at the indicated M.O.I. (**A**) Cells were fixed at 24 h post infection (hpi) and infected cells were visualized by indirect immunofluorescence (IF) staining of HCoV-OC43 N protein (green). Cell nuclei were visualized by DAPI staining. (**B**) HCoV-OC43 NP, IFITMs and β-actin were determined by Western blot assays. (**C**) Intracellular viral RNA was quantified by qRT-PCR assay and presented as copies per 100 ng total RNA. Error bars indicate standard deviations (n = 4). (D) Viral yields were determined with a plaque assay. Error bars indicate standard deviations (n = 4). ** indicates *p* <0.001 (student *t* test).

**Fig. 2.**
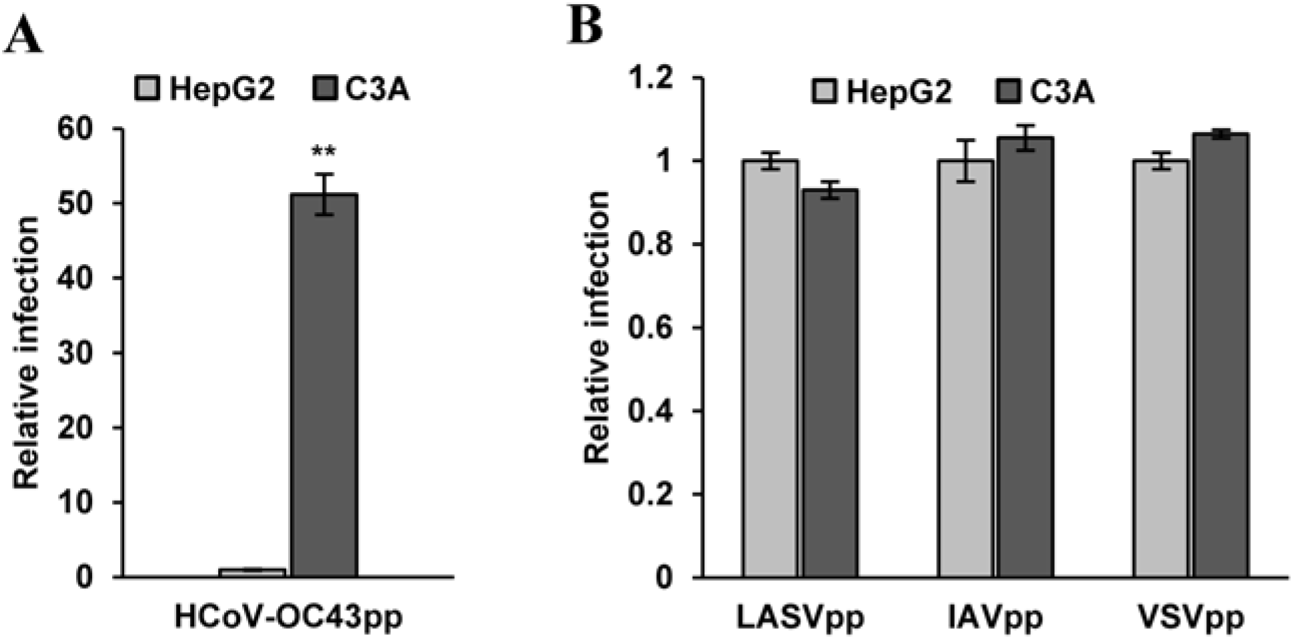
C3A cells support more efficient entry of lentiviral particles pseudotyped with HCoV-OC43 envelope proteins than HepG2 cells. HepG2 and C3A cells were infected with HCoV-OC43pp (A), IAVpp, VSVpp or LASVpp (B). Luciferase activities were determined at 72 hpi. Relative infection represents the luciferase activity from C3A normalized to that of HepG2 cells. Error bars indicate standard deviations (n = 6). ** indicates *p* <0.001 (student *t* test).

### IFITM proteins modulate HCoV-OC43 infection of C3A and HepG2 cells in a similar extent

We reported previously that IFITM proteins differentially modulate HCoV-OC43 entry into target cells. While IFITM1 inhibits the virus entry, IFITM2 and IFITM3 enhance the cellular entry of this virus (42). To investigate whether the differential expression of IFITM proteins in the two cell lines is responsible for their difference in HCoV-OC43 infection efficiency, we examined IFITM protein expression by Western blot assays and found the two hepatoma cell lines expressed similar levels of IFITM1 and IFITM2/3 (Fig. 1B). Because the C-terminal variable regions of IFITM1 and IFITM3 control the inhibition and enhancement of HCoV-OC43 entry (42), respectively, we further compared the effects of IFITM1 and C-terminal region exchanged IFITM proteins on the virus infection. As shown in Fig. 3, in spite of their distinct susceptibility, expression of IFITM1-EX2, a mutant IFITM1 protein with its C-terminal domain replaced with the C-terminal domain of IFITM3 (42), and IFITM3-EX2, a mutant IFITM3 protein with its C-terminal domain replaced with the C-terminal domain of IFITM1 (42), significantly enhanced and inhibited HCoV-OC43 infection of both cell lines, respectively, as evidenced by the significant changes in infected cell percentage (Fig. 3A), reduced viral nucleocapsid protein expression (Fig. 3B), intracellular RNA accumulation (Fig. 3C) and yields of progeny virus production (Fig. 3D). Moreover, pseudotyped lentiviral infection assay further demonstrated that IFITM1, IFITM1-EX2 and IFITM3-EX2 modulated HCoV-OC43 envelope proteins mediated entry in a similar extent in the two cell lines (Fig. 3E). Accordingly, we concluded that IFITM proteins were not responsible for the observed differential susceptibility of the two hepatoma cell lines to HCoV-OC43 infection.

**Fig. 3.**
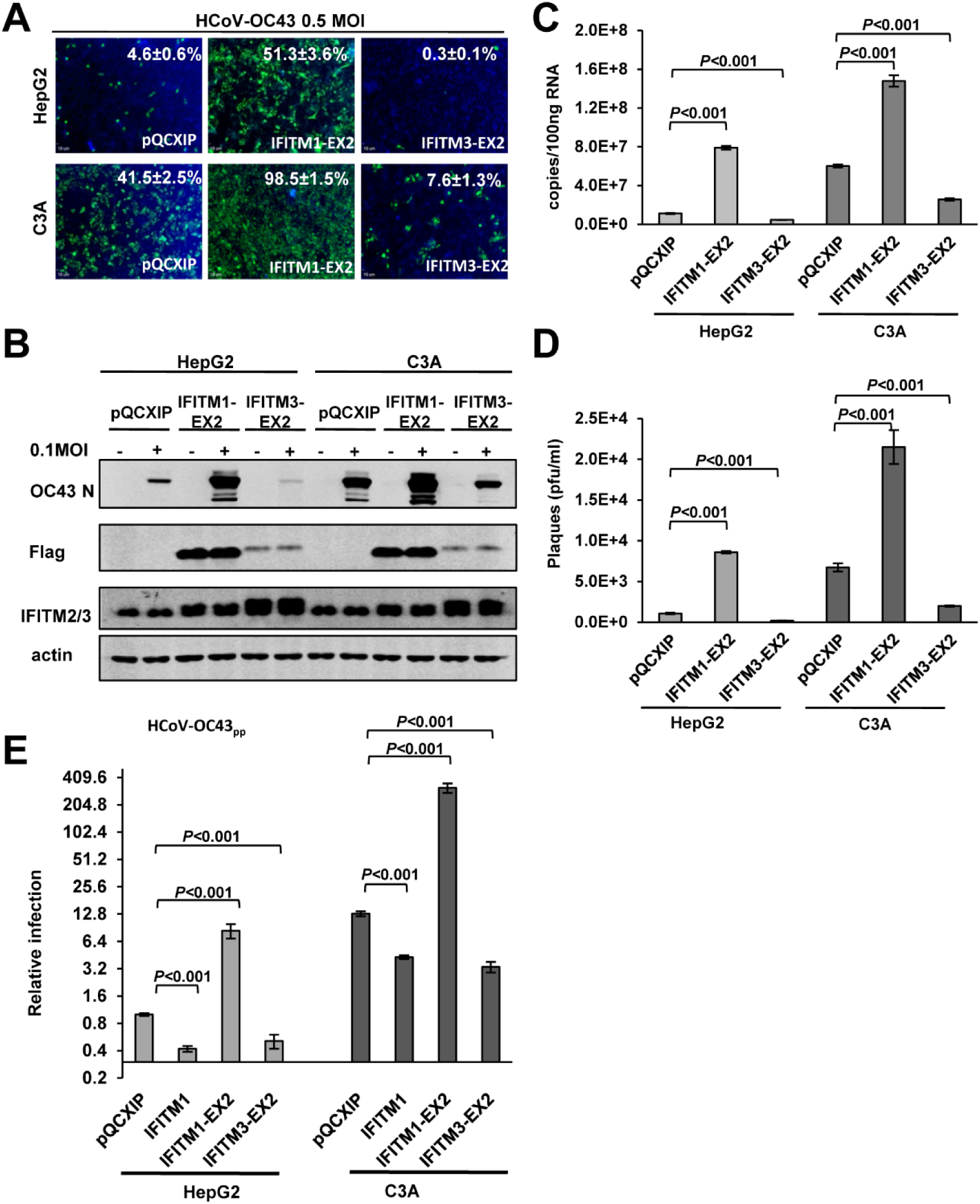
IFITMs modulate HCoV-OC43 infection of HepG2 and C3A cells in a similar extent and *via* the same mechanism. HepG2 and C3A were stably transduced with a control retroviral vector (pQCXIP) or a retroviral vector expressing a N-terminally flag tagged IFITM1-EX2 or IFITM3-EX2. The resulting cell lines were infected with HCoV-OC43 at 0.1 MOI. (**A**) Cells were fixed at 24 hpi and virally infected cells were visualized by IF staining of HCoV-OC43 N protein (green). Cell nuclei were visualized by DAPI staining. (**B**) HCoV-OC43 NP and IFITM were determined by Western blot assays. β-actin served as a loading control. (**C**) Intracellular viral RNA was quantified by a qRT-PCR assay and presented as copies per 100 ng total RNA. Error bars indicate standard deviations (n = 4). (**D**) Viral yields were determined with a plaque assay. Error bars indicate standard deviations (n = 4). (E) HepG2 and C3A stably transduced with a control retroviral vector (pQCXIP) or a retroviral vector expressing IFITM1, IFITM1-EX2 or IFITM3-EX2 were infected with HCoV-OC43pp. Luciferase activities were determined at 72 hpi. Relative infection represents the luciferase activity normalized to that of HepG2 cells transduced with empty vector (pQCXIP). Error bars indicate standard deviations (n = 6).

### LY6E inhibits the entry mediated by human CoV envelope spike proteins and is responsible for the differential susceptibility of C3A and HepG2 cells to HCoV-OC43 infection

In order to identify host cellular proteins that may enhance HCoV-OC43 infection of C3A cells or suppress the virus entry into HepG2 cells, we first compared the expression of several cellular genes with known activity to restrict or enhance virus entry into target cells. As shown in Fig. 4A, we found that ADAP2, GILT and LY6E mRNA expressed at significantly higher levels in HepG2 cells. While the expression of ADAP2 and GILT did not inhibit HCoV-OC43pp infection (Fig. 4B), expression of LY6E in Flp-In TREx 293 cells efficiently suppressed the infection of lentiviral particles pseudotyped with the envelope glycoproteins of all the human CoVs including SARS-CoV-2, except for SARS-CoV (Fig. 4C and 4D). In addition, LY6E enhanced the infection of IAVpp (Fig. 4D). In agreement with our previous report, expression of GILT inhibited SARS-CoVpp infection (38). To further confirm the role of LY6E in HCoV-OC43 infection, we showed that ectopic expression of LY6E in C3A and A549 cells significantly reduced their susceptibility to the virus infection, whereas reducing the expression of LY6E in HepG2 cells by shRNA knockdown significantly increased HCoV-OC43 infection (Fig. 5). The results presented above imply that LY6E is a restriction factor for human CoVs and responsible for the differential susceptibility of C3A and HepG2 cells to HCoV-OC43 infection.

**Fig. 4.**
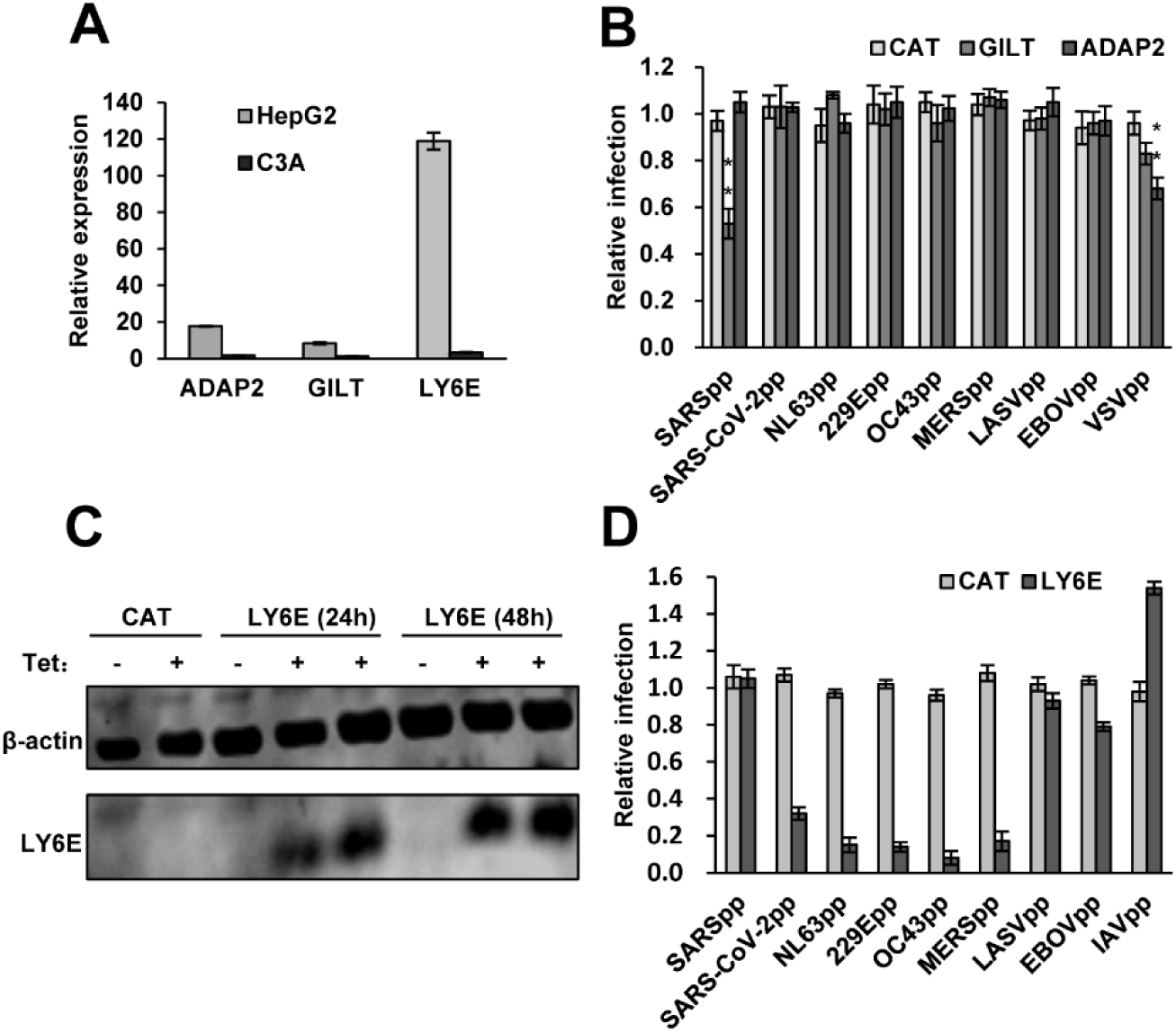
LY6E efficiently suppresses human coronavirus spike protein-mediated entry. (A) Levels of Ly6E, GILT and ADAP2 mRNA expression in HepG2 and C3A cells were determined by qRT-PCR assays and normalized to the level of GAPDH. (B) Flp-In T-Rex 293-derived cell lines expressing control protein CAT, GILT or ADAP2 were cultured in the absence or presence of tet for 24 h. The cells were infected with HCoV-OC43pp and other indicated pseudoviral particles and intracellular luciferase activity were determined at 48 hpi. Relative infection is the ratio of luciferase activity in the same cells cultured in the presence of tet over that in the absence of tet. The error bars refer to standard deviations (n=4). (C) Flp-In T-Rex 293-derived cell line expressing a control protein CAT or LY6E were cultured in the absence or presence of tet. Cells were harvested at the indicated time after the addition of tet. Intracellular expression of LY6E was detected by a Western blot assay. β-actin served as a loading control. (D) Flp-In T-Rex 293-derived cell lines expressing LY6E were cultured in the absence or presence of tet for 24 h. The cells were then infected with lentiviral particles pseudotyped with the envelope protein of the indicated viruses. Luciferase activities were determined at 48 hpi. Relative infection is the ratio of luciferase activity in the same cells cultured in the presence of tet over that in the absence of tet. The error bars refer to standard deviations (n=4). **, *p* <0.001 compared to the control cells expressing CAT.

**Fig. 5.**
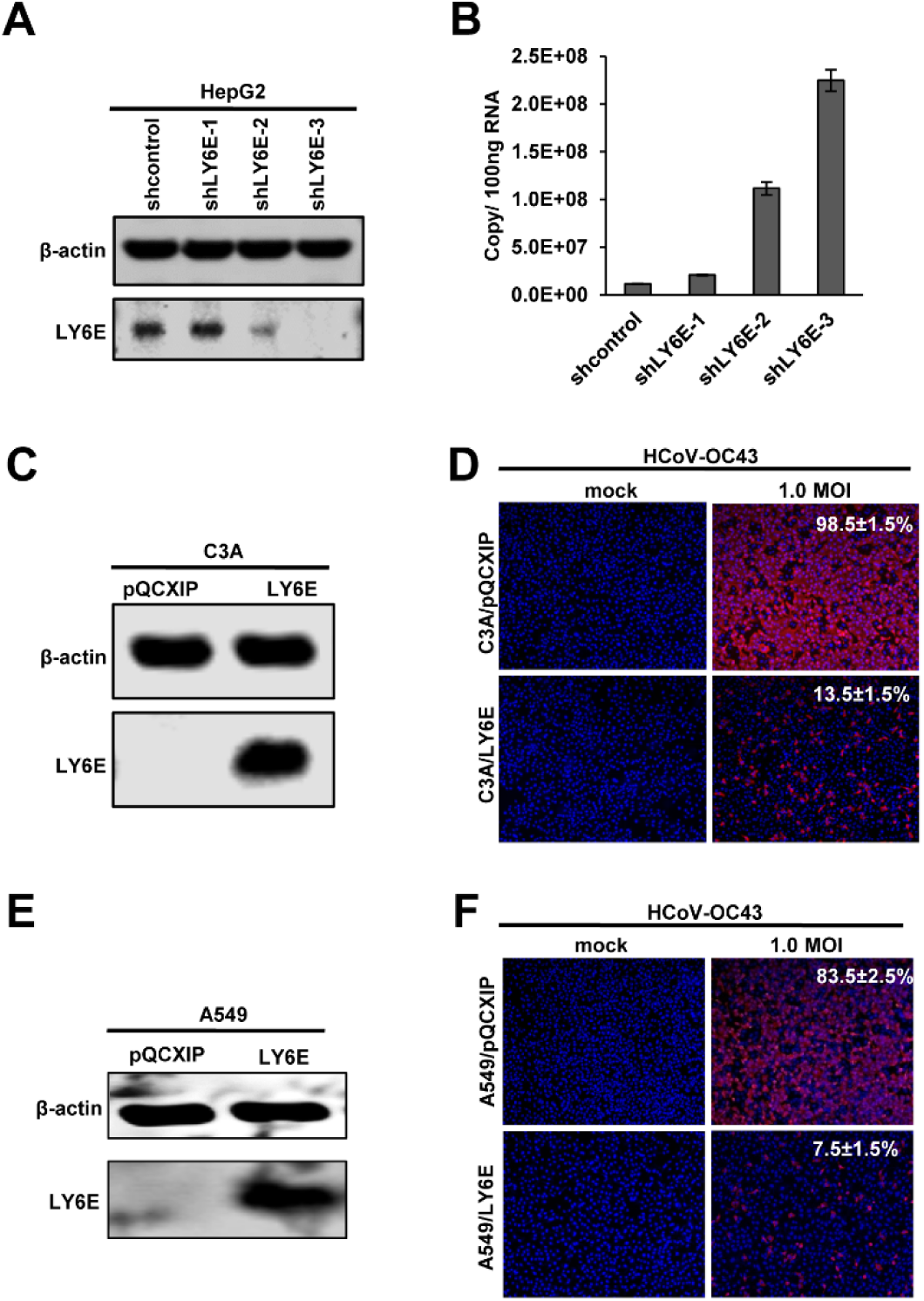
LY6E inhibits HCoV-OC43 infection in human hepatoma (HepG2 and C3A) and lung cancer (A549) cells. (A) HepG2 cells were stably transduced with scramble shRNA or shRNA targeting LY6E mRNA. The level of intracellular LY6E expression was determined by Western blot using a rabbit polyclonal antibody against LY6E. β-actin served as a loading control. (B) HepG2 cells stably expressing the scramble shRNA or LY6E specific shRNA were infected with HCoV-OC43 at an MOI of 1.0. Cells were harvested at 24 hpi and intracellular viral RNA was quantified by qRT-PCR assay and presented as copies per 100 ng total RNA. Error bars indicate standard deviations (n = 4). (C to F) C3A or A549 cells were stably transduced with an empty retroviral vector (pQCXIP) or retroviral vector expressing LY6E and infected with HCoV-OC43 at the indicated MOI. The expression of LY6E in the cell lines was confirmed by a western blot assay. β-actin served as a loading control (C and E). The cells were fixed at 24 hpi. The infected cells were visualized by IF staining of HCoV-OC43 N protein (red). Cell nuclei were visualized by DAPI staining (D and F).

### LY6E restriction of human coronavirus entry depends on GPI-anchor and the evolutionally conserved L36 residue

LY6E is a member of the LY6/uPAR superfamily. Like most LY6 family members, LY6E contains ten cysteines that form a highly conserved, three-finger folding motif through disulfide bonding and localizes on the plasma membrane of cells *via* glycosylphosphatidylinositol (GPI) anchoring. LY6E is ubiquitously expressed in many cell types and functions in modulation of cell signal transduction (41). Recent studies revealed that human LY6E promotes the entry of HIV (46, 47) and multiple enveloped RNA viruses from several viral families (48). Moreover, the enhancement of RNA viral infection is a conserved function of all the mammalian LY6E orthologs examined thus far. Particularly, substitution of the evolutionally conserved residue L36 with alanine (A) completely abolished the viral enhancement activity of LY6E (48). Interestingly, we found that L36A substitution also abolished the activity of LY6E to restrict the entry of human CoVs (Fig. 6). As anticipated, N99A substitution that disrupts the addition of GPI anchor also abrogated the inhibitory effects of LY6E on human CoV entry (Fig. 6). These results indicate that proper interaction of LY6E with other viral/cellular components *via* the conserved residue L36 and localization in the specific cell membrane microdomians are required for LY6E restriction of human CoV entry.

**Fig. 6.**
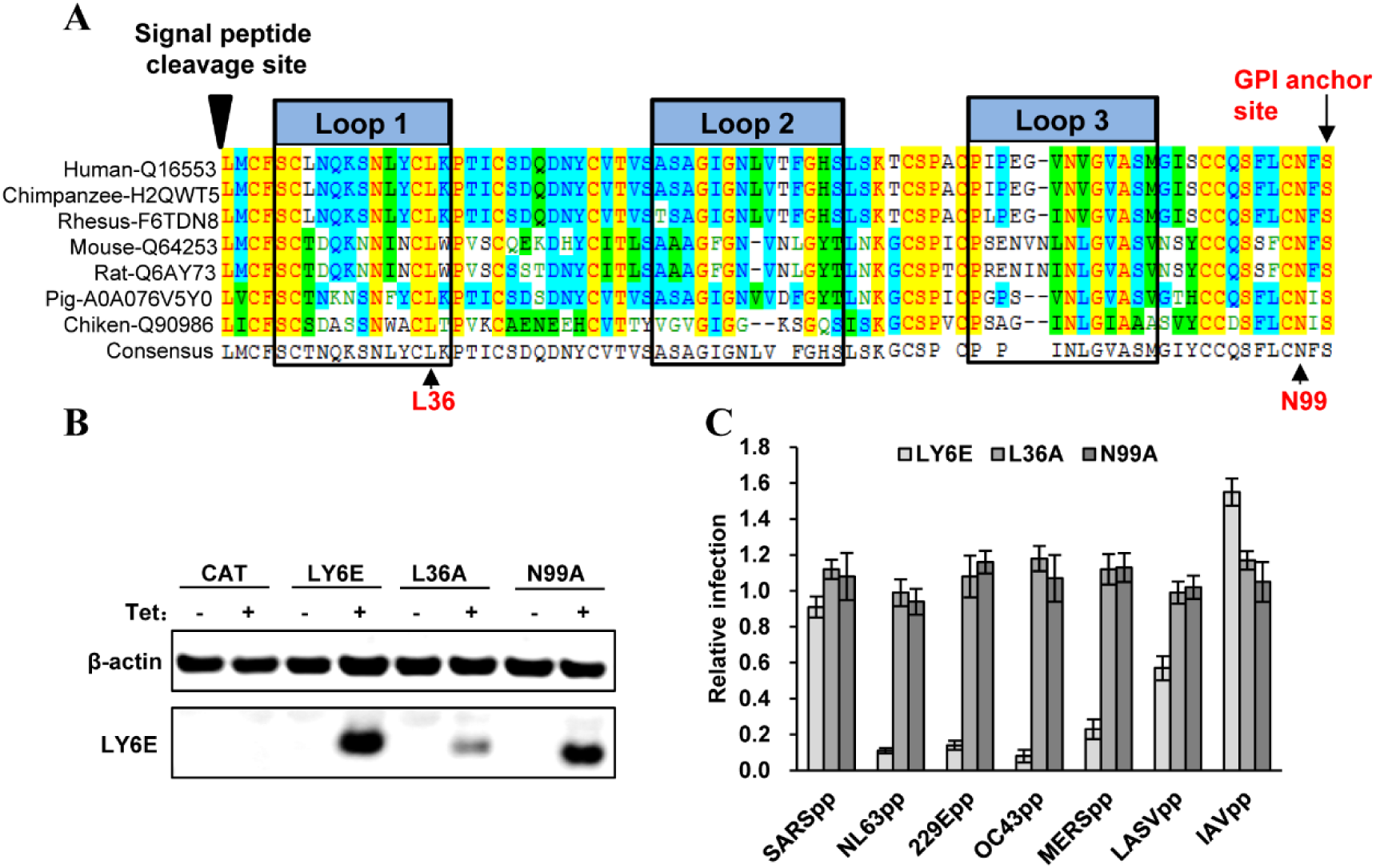
Identification of critical structure motifs essential for LY6E to restrict human coronavirus entry. (A) The amino acid sequence alignment of LY6E from multiple vertebrate species is conducted and “three finger-fold” structure is highlighted with black box. The conserved L36 as well as GPI anchor and N99 glycosylation sites are indicated (B) Flp-In T-Rex 293-derived cell lines expressing a control protein CAT, wild-type or mutant LY6E were cultured in the absence or presence of tet for 24 h. Intracellular LY6E expression were detected by a Western blot assay. β-actin served as a loading control. (C) Flp-In T-Rex 293-derived cell lines expressing the wild-type or mutant LY6E were cultured in the absence or presence of tet for 24 h. The cells were then infected with the indicated pseudotyped lentivirus. Luciferase activities were measured at 48hpi. Relative infection is the ratio of luciferase activity in the same cells cultured in the presence of tet over that in the absence of tet. The error bars refer to standard deviations (n=4).

### Activation of CoV entry by TMPRSS2 expression fails to evade LY6E restriction of CoV entry

It was reported by others and us that expression of cell membrane associated serine protease TMPRSS2 enhances SARS-CoV and SARS-like bat CoV entry (32-34). More importantly, the TMPRSS2-enhanced entry can evade IFITM3 restriction (*Mei Zheng, et al, manuscript under review*), presumably because the cellular protease activates the viral fusion at cell surface or early endosomes where IFITM3 expression at a relatively lower levels and thus fails to inhibit viral fusion. To determine the effects of TMPRSS2 expression on LY6E restriction of human CoV entry, Flp-In TREx 293-derived cell line expressing LY6E were transfected with a control vector (pCAGGS) or a plasmid expressing human TMPRSS2 and cultured in the absence or presence of tetracycline (tet) for 24 h. The cells were then infected with the indicated pseudotyped lentiviruses. As shown in Fig. 7A, in absence of tet to induce LY6E, expression of TMPRSS2 significantly enhanced the infection of SARS-CoVpp, MERS-CoVpp and HCoV-229Epp, but not the infection of other human CoVpp and LASVpp. Interestingly, LY6E significantly inhibited the infection of MERS-CoVpp and HCoV-229Epp in the cells without or with ectopic expression of TMPRSS2. Moreover, LY6E failed to inhibit SARS-CoVpp infection in the absence of TMPRSS2 expression, but significantly inhibited SARS-CoVpp infection in the cells expressing TMPRSS2 (Fig. 7B). Therefore, unlikely IFITM3, expression of TMPRSS2 cannot evade LY6E restriction of human CoV entry. Instead, LY6E restriction of the entry of human CoVs, particularly SARS-CoV, is regulated by the expression of cellular proteases that previously known to activate the fusion activity of viral spike proteins.

**Fig. 7.**
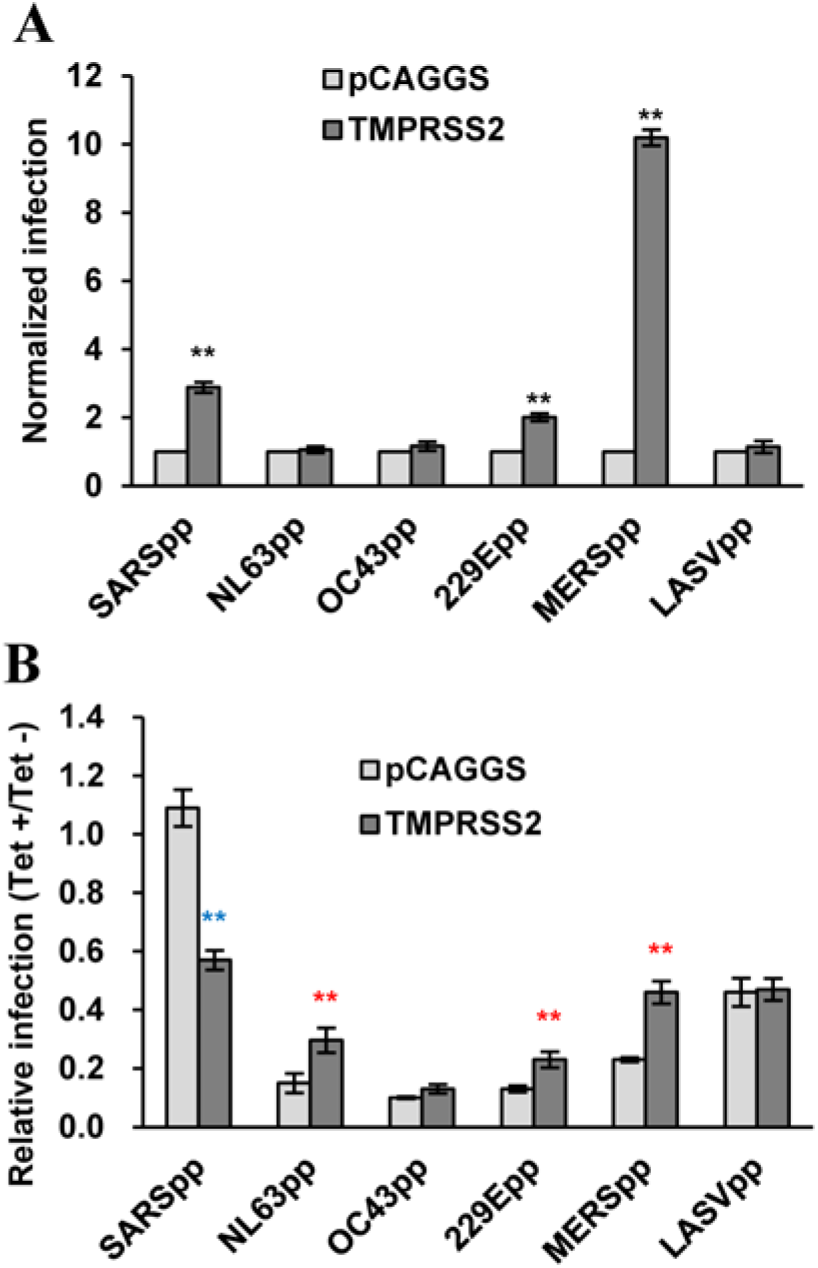
LY6E inhibits TMPRSS2 enhanced entry of human coronaviruses. Flp-In T-Rex 293-derived cell line expressing LY6E were transfected with a control vector (pCAGGS) or a plasmid expressing human TMPRSS2 and cultured in the absence or presence of tet for 24 h. The cells were then infected with the indicated pseudotyped lentivirus. Luciferase activities were measured at 48hpi. (**A**) The effect of TMPRSS2 expression on pseudotyped virus infection is normalized to infection efficiency of the cells transfected with control vector plasmid (set as 1). Error bars indicate the standard deviation (n=4). (**B**) Relative infection refers to the ratio of the luciferase activity in the cells cultured in the presence of tet over that in the cells cultured in the absence of tet. Error bars indicate the standard deviation (n=4). **, *p* <0.001, comparing to cells transfected with pCAGGS vector.

### Amphotericin B treatment does not compromise LY6E restriction of human CoV entry

Amphotericin B (AmphoB) is antifungal medicine that binds with ergosterol in fungal cell membranes, forming pores that cause rapid leakage of monovalent ions and subsequent fungal cell death. AmphoB can also bind to cholesterol in mammalian cell membrane, albeit at a lesser affinity than to fungal ergosterol (49). The cholesterol-enriched plasma membrane microdomains known as lipid rafts play important roles in the entry and egress of many enveloped viruses (50, 51). Particularly, AmphoB treatment had been shown to significantly compromise IFITM restriction of IAV entry (52) and attenuate IFITM enhancement of HCoV-OC43 infection (42). In this study, we further demonstrated that AmphoB treatment also efficiently attenuated the restriction of IFITM3 on the infection of SARS-CoVpp, MERS-CoVpp, HCoV-NL63pp, HCoV-229Epp and IAVpp, but not LASVpp (Fig. 8A). However, AmphoB treatment altered neither the restriction activity of LY6E on the infection of human CoV spike protein-pseudotyped lentiviruses nor the enhancement of LY6E on IAVpp infection (Fig. 8B). These results strongly imply that LY6E modulates virus entry *via* a distinct mechanism.

**Fig. 8.**
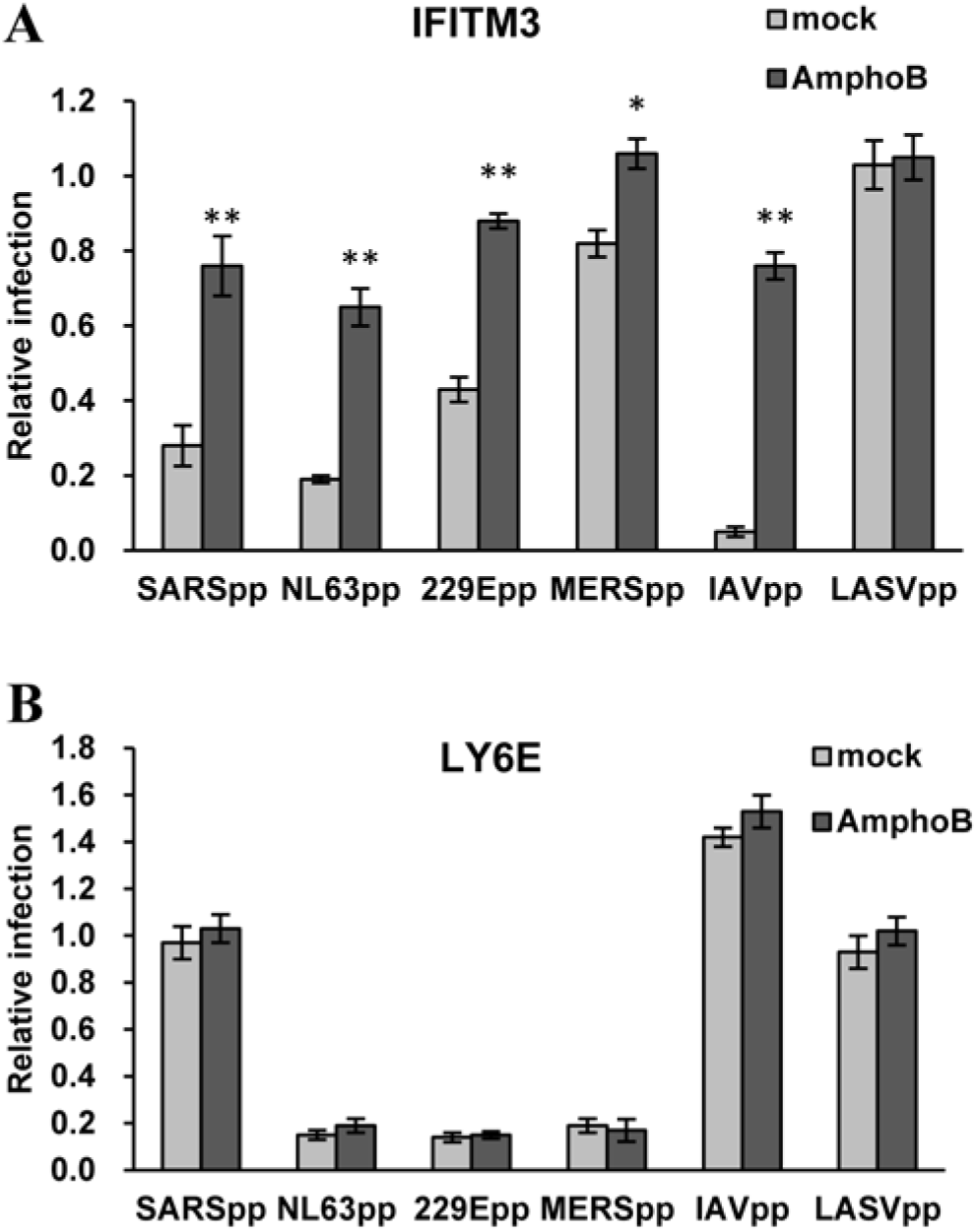
Amphotericin B treatment compromises IFITM3 inhibition of human coronavirus entry, but have no impact on Ly6E inhibition of human coronavirus entry. Flp-In T-Rex 293-derived cell line expressing IFITM3 (**A**) or LY6E (**B**) were cultured in the absence or presence of tet for 24 h. The cells were then infected with the indicated pseudotyped lentivirus in the presence or absence of 1μM AmphoB. Luciferase activity was measured at 48 hr post-infection. Relative infection is the ratio of luciferase activity in the same cells cultured in the presence of tet over that in the absence of tet. The error bars refer to standard deviations (n=4). **, *p* <0.001, compared to mock treatment.

## DISCUSSION

LY6E was initially identified as a cell surface marker to discriminate immature from mature thymocytes subsets (53). The primary function of LY6E has been associated with immune regulation, specifically in modulating T cell activation, proliferation, development (54). In addition to lymphocytes, LY6E mRNA can also be detected in liver, spleen, uterus, ovary, lung, and brain and its expression can be induced by type I IFN in a cell-type specific manner (53). However, LY6E is not a typical antiviral effector protein. Instead, LY6E was reported to promote the infection of enveloped RNA viruses from several viral families (48) and modulates HIV-1 infection in a manner dependent on the level of CD4 expression in target cells (46, 47). Our finding that LY6E restricts human CoV infection and characterization of its antiviral effects shed new lights on the mode of LY6E action on virus entry in general.

First, either the enhancement or restriction of LY6E on virus entry depends on its GPI anchor (Fig. 6) (46-48). GPI-anchored proteins are preferentially located in lipid rafts, the plasma membrane microdomains enriched in glycosphingolipids and cholesterol as well as protein receptors or ligands. Lipid rafts are considered to compartmentalize membrane processes by facilitating the interaction of protein receptors and their ligands/effectors to modulate membrane functions, such as signal transduction, membrane fusion, vesicle budding and trafficking. Lipid rafts also involve in the entry and egress of many viruses. For instance, both HIV-1 receptor (CD4) and coreceptors are localized in lipid rafts. Yu and colleagues elegantly demonstrated recently that LY6E enhances HIV-1 infection of CD4+ T cells and monocytic THP1-cells by promoting the expansion of viral fusion pore induced by HIV-1 Env (47). Furthermore, LY6E was found to be the receptor of mouse endogenous retroviral envelope Syncytin-A and interaction of LY6E with Syncytin-A induces the syncytiotrophoblast fusion and placental morphogenesis (55, 56). However, Mar and colleagues showed that LY6E enhances IAV infection of cells by promoting a viral replication step after viral nucleocapsid escape from endosomes, but before viral RNP nuclear translocation, *i*.*e*., most likely the uncoating of nucleocapsids (48). Interestingly, the results presented in Fig. 4 indicate that LY6E significantly enhanced the infection of IAVpp, suggesting that the LY6E enhancement of IAV infection is, at least in part, through promoting the entry into target cells, possibly also by enhancing viral fusion. Considering the broad inhibitory effects of LY6E on human CoVs and its fusogenic or fusion-modulating activity, we speculate that LY6E might inhibit the membrane fusion triggered by CoV spike proteins. However, the role of LY6E on endocytosis and endocytic vesicle trafficking cannot be ruled out. These hypotheses are currently under investigation.

Second, in addition to GPI anchor, the evolutionally conserved amino acid residue L36 is also required for both the enhancement and restriction of virus entry into target cells by LY6E (Fig. 6) (48). It can be speculated that this specific residue may mediate an interaction with other cellular membrane proteins to module viral entry. The fact that LY6E enhances viral infectivity in a cell type-specific manner, with the strongest phenotype in cells of fibroblast and monocytic lineages (48), does indicate the involvement of other host cellular factors. Variations in the abundance of expression, as well as the localization of LY6E and its associated proteins or lipids, may explain the differential effects of LY6E on the infection of different viruses in different cell types (Fig. 4 and 7). However, LY6E enhancement of RNA virus infection appears to be independent of type I interferon response and other ISG expression (48). Particularly, enhancement of viral infection in Huh7.5 cells that do not have basal levels of IFITM protein expression indicates that LY6E enhancement of RNA viral infection is most likely not through modulating the function of IFITM proteins (42). This notion is further supported by the finding that LY6E and Syncitin-A mediated syncytiotrophoblast fusion can be inhibited by IFITM proteins (57, 58).

Third, studying the effects of LY6E on HIV-1 infection of CD4 low-expressing cells, such as Jurkat T cells and primary monocyte-derived macrophages, revealed that HIV-1 entry was inhibited by LY6E (46). This appears due to the LY6E-induced reduction of lipid raft-associated CD4 on the surface of these cells. It was demonstrated that LY6E can promote CD4 endocytosis and mobilize lipid raft-associated CD4 molecules to non-raft microdomains. Such a receptor down-regulation significantly reduced HIV-1 binding and infection of CD4 low-expressing cells (macrophages), but did not significantly impact the binding of HIV-1 to CD4 high-expressing cells, which allows for LY6E to predominantly enhance HIV-1 infection of CD4+ T lymphocytes by promotion of membrane fusion (46). It is, therefore, possible that LY6E inhibition of human CoV infection is due to the down-regulation of lipid raft-associated CoV receptors. However, the differential effects of LY6E on the infection of SARS-CoVpp, SARS-CoV-2pp and HCoV-NL63pp, that share the ACE2 receptor, does not support such a hypothesis (Fig. 4).

Finally, the findings that LY6E inhibits human CoV entry cannot be evaded by ectopic expression of membrane-associated serine protease TMPRSS2 and compromised by AmphoB treatment strongly indicate that LY6E modulates virus entry *via* a distinct mechanism from that IFITM proteins do (Figs. 7 and 8). Specifically, inhibition of TMPESS2-enhanced CoV entry implies that LY6E most likely blocks virus entry at plasma membrane or in early endosomes. Moreover, IFITMs impede viral fusion by decreasing membrane fluidity and curvature (37). AmphoB can bind cholesterol in cell membranes to increase membrane fluidity and planarity and consequentially rescue IFITM inhibition of virus entry (52). Interestingly, AmphoB only neutralize the antiviral effects of IFITM2 and IFITM3, but has little effect on IFITM1 restriction of virus entry (52). While IFITM1 is predominantly located in the plasma membrane or early endosomes, IFITM2 and 3 are mainly localized in the later endosomes and lysosomes. Due to their differential subcellular localization, IFITM1 mainly restricts the viruses that enter the cells at cell surface or in the early endosomes, such as parainfluenza viruses and hepatitis C virus (59, 60), IFITM2 and 3 primarily restrict the infection of viruses that enter the cells at later endosomes and/or lysosomes (43, 61, 62). Because AmphoB is endocytosed quite rapidly leading to its concentration in the late endosomes and lysosomes, it more efficiently alleviates the effect of IFITM2 and 3, but not IFITM1, on virus entry (52). Similarly, the failure of AmphoB to attenuate the antiviral effects of LY6E against human CoVs is most likely due to its predominant cell surface localization and inhibition of an early step of CoV entry.

In summary, while it is very interesting to know that LY6E is capable of modulating the entry of many RNA viruses, we only begin to uncover the mechanism of this fascinating host factor and define its pathobiological role in virus infection (41, 63). Further understanding the role and mechanism of LY6E in viral infections will establish a scientific basis for development of therapeutics to harness its function for the treatment of viral diseases.

## MATERIALS AND METHODS

### Cell culture

Human hepatoma cell lines HepG2 and C3A, a sub-clone of HepG2 (ATCC HB-8065) were purchased from ATCC and cultured in DMEM/F12 medium supplemented with 10% heat-inactivated fetal bovine serum (FBS) (Invitrogen). Lung cancer cell line A549 were obtained from ATCC and maintained in DMEM supplemented with 10% FBS. GP2-293 and Lenti-X 293T cell Lines were purchased from Clontech and cultured in DMEM supplemented with 10% FBS and 1 mM Sodium pyruvate (Invitrogen). Flp-In TREx 293 cells were purchased from Invitrogen and maintained in DMEM supplemented with 10% FBS, 10 μg/ml blasticidin (Invitrogen) and 100 µg/ml Zeocin (Invivogen) (64). Flp-In TREx 293-derived cell lines expressing LY6E, GILT, ADAP2, or IFITM3 were cultured in DMEM supplemented with 10% FBS, 5 μg/ml blasticidin and 250 μg/ml hygromycin.

### Viruses

HCoV-OC43 (strain VR1558) were purchased from ATCC and amplified in HCT-8 cells according to the instruction from ATCC. Virus titers were determined by a plaque assay as described previously (42).

### Antibodies

Monoclonal antibody against FLAG tag (ANTI-FLAG M2) and β-Actin were purchased from Sigma (Cat.No. F1804 and A2228, respectively). Monoclonal antibody against human IFITM1 (Cat.No. 60047-1), rabbit polyclonal antibody against human IFITM3 (Cat.No. 11714-1-AP), which also efficiently recognizes IFITM2 and weakly cross-reacts with IFITM1, were purchased from Proteintech Group, Inc. Mouse monoclonal antibody against HCoV-OC43 nucleocapsid (NP) protein was purchased from Millipore (Cat.No. MAB9012). Rabbit polyclonal antibody against human LY6E was obtained from proteintech (Cat.No. 22144-1-AP).

### Plasmid construction

The cDNA molecules of ADAP2 and LY6E were purchased from OriGene (Cat. No. RC207501 and RC211373, respectively) and cloned into pcDNA5/FRT-derived vector as described previously (42). Ly6E and N-terminally FLAG-tagged human IFITM1, IFITM3 and their mutants were cloned into pQCXIP vector (Clontech) between the NotI and BamHI sites as previously described (42, 43). pcDNA5/FRT-derived plasmids expressing chloramphenicol acetyltransferase (CAT), N-terminally FLAG-tagged human IFITM3 were reported previously (64-66).

Plasmids expressing HCoV-OC43 spike (S) and HE proteins, VSV G protein, H1N1 IAV (A/WSN/33) hemagglutinin (HA) and neuraminidase (NA), LASV GP protein, murine leukemia virus (MLV) envelope protein, HCoV-NL63, HCoV-229E, SARS-CoV and MERS-CoV spike protein were described previously (67, 68). The codon-optimized (for human cells) SARS-CoV-2 spike gene, which is based on NCBI Reference Sequence YP_009724390.1, was purchased from GeneScript and cloned into pCAGGS vector as described previously (69). pRS-derived retroviral vectors expressing a scramble shRNA and shRNA targeting the mRNA of human LY6E were obtained from OriGene (Cat. No. TR311641).

Plasmid pNL4-3.Luc.R^-^E^-^ was obtained through the NIH AIDS Research and Reference Reagent Program (70, 71). Angiotensin I converting enzyme 2 (ACE2), aminopeptidase N (APN), and dipeptidyl peptidase-4 (DPP4) cDNA clones were obtained from Origene, and cloned into a pcDNA3 vector (Invitrogen) to yield plasmid pcDNA3 /ACE2, pcDNA3 /APN and pcDNA3/DDP4, respectively (69).

### Package of pseudotyped retroviral particles

The various viral envelope protein pseudotyped lentiviruses bearing luciferase reporter gene as well as VSV G protein pseudotyped Moloney murine leukemia virus (MMLV)-derived retroviral vectors (pQCXIP) expressing wild-type and mutant human IFITM, LY6E, or pRS vector-derived plasmid expressing a scrambled shRNA or shRNA specifically targeting human LY6E were packaged as reported previously. Each pseudotype was titrated by infection of cells with a serial dilution of pseudotype preparations. The modulation of IFITM or LY6E on the transduction of a given pseudotype was determined with a titrated amount of pseudotypes that yield luciferase signal between 10,000 to 1,000,000 light units per well of 96-well plates (69, 72). For a given pseudotype, the input of pseudoviral particles is consistent across all the experiments.

### Establishment of cell lines stably expressing wild-type and mutant IFITM or LY6E proteins or shRNA

HepG2, C3A or A549 cells in each well of 6-well plates were incubated with 2 ml of Opti-MEM medium containing pseudotyped retroviruses and centrifuged at 20 °C for 30 minutes at 4,000×g. Forty-eight hours post transduction, cells were cultured with media containing 2 µg/ml of puromycin for two weeks. The antibiotic resistant cells were pooled and expanded into cell lines stably expressing wild-type or mutant IFITM or LY6E proteins or shRNA targeting LY6E. Flp-In TREx 293-derived cell lines expressing IFITM, GILT, ADAP2 or LY6E proteins in a tetracycline (tet) inducible manner were established as previously described (64, 66).

### Immunofluorescence

To visualize HCoV-OC43 infected cells, the infected cultures were fixed with 2% paraformaldehyde for 10 min. After permeabilization with 0.1% Triton X-100, the cells were stained with a monoclonal antibody (541-8F) recognizing HCoV-OC43 NP protein. The bound antibodies were visualized by using Alexa Fluor 488-labeled (green) goat anti-mouse IgG or Alexa Fluor 555-labeled (red) goat anti-mouse IgG, Cell nuclei were counterstained with DAPI.

### Western blot assay

Cells were lysed with 1× Laemmli buffer. An aliquot of cell lysate was separated on NuPAGE® Novex 4-12% Bis-Tris Gel (Invitrogen) and electrophoretically transferred onto a nitrocellulose membrane (Invitrogen). The membranes were blocked with PBS containing 5% nonfat dry milk and probed with the desired antibody. The bound antibodies were visualized with IRDye secondary antibodies and imaging with LI-COR Odyssey system (LI-COR Biotechnology).

### Real-time RT-PCR

HCoV-OC43 RNA was quantified by a qRT-PCR assay described previously (42). To determine the level of ISG mRNA, total cellular RNA was extracted using TRIzol reagent (Invitrogen) and the same amount of total cellular RNA was reverse-transcribed with SuperScript III kit ((Invitrogen). Quantitative RT-PCR was performed using iTaq universal SYBR Green Supermix (Bio-Rad) with the following primers: LY6E, 5’-GTACTGCCTGAAGCCGACCATC-3’ and 5’-AGATTCCCAATGCCGGCACTAG-3’; ADAP2, 5’-AGCTGTCATCAGCATTAAG-3’ and 5’-ACTATCTCCTTCCCACTTTC-3’; GILT, 5’-AATGTGACCCTCTACTATGAAG-3’ and 5’-ACGCTGGTGCCCTACGGAAACG-3’; GAPDH, 5’-GAAGGTGAAGGTCGGAGTCAAC-3’ and 5’-CAGAGTTAAAAGCAGCCCTGGT-3’. Gene expression was calculated using the 2^-ΔΔCT^ method, normalized to GAPDH as described previously (31, 40).

### Luciferase assay

Flp-In TREx 293**-**derived IFITM-expressing cell lines were seeded into 96-well plates with black wall and clear bottom and transfected with an empty vector plasmid or plasmids encoding ACE2, APN, or DPP4 to express viral receptors. For Huh7.5-derived IFITM-expressing cell lines, cells were seeded into black wall 96-well plates. Cells were infected at 24 h post transfection or infected with desired pseudotyped lentiviral particles for 2 h, and then replenished with fresh media. Two days post infection, the media were removed and cells were lysed with 20 µl/well of cell lysis buffer (Promega) for 15 min, followed by adding 50 µl/well of luciferase substrate (Promega). The firefly luciferase activities were measured by luminometry in a TopCounter (Perkin Elmer) (69).

### Statistical analyses

All the experiments were repeated at least three times. Differences between control sample and tests were statistically analyzed using Student’s *t* tests or one-way analysis of variance (ANOVA). *p*-values less than 0.05 were considered statistically significant.

## Funding Information

This work was supported by grants from the National Institutes of Health, USA (AI113267) to J.-T. Guo, National Natural Science Foundation of China (81772173 and 81971916) and National Science and Technology Mega-Project of China (2018ZX10301-408-002) to X. Zhao and The Commonwealth of Pennsylvania through the Hepatitis B Foundation.

